# Neuronal function and dopamine signaling evolve at high temperature in *Drosophila*

**DOI:** 10.1101/585422

**Authors:** Jakšić Ana Marija, Karner Julia, Nolte Viola, Hsu Sheng-Kai, Barghi Neda, Mallard François, Otte Kathrin Anna, Svečnjak Lidija, Senti Kirsten-André, Schlötterer Christian

**Affiliations:** Institut für Populationsgenetik, Vetmeduni Vienna, Vienna, Austria; Vienna Graduate School of Population Genetics, Vetmeduni Vienna, Vienna, Austria; University of Zagreb, Faculty of Agriculture, Zagreb, Croatia

**Author notes:** Department for Molecular Biology and Genetics, Cornell University, NY, USA. Institut de Biologie de l’École Normale Supérieure, Paris, France (Mallard François).

## Introduction

Adaptation to local environments is a general principle in nature. In the light of the ongoing climatic change, the mechanisms for temperature adaptation are of particular interest^1–2^. Many essential biological processes are temperature sensitive, and neuronal activity^3–7^ is a particularly prominent one as it affects essential, fitness-related traits such as behavior and locomotion^8^. Here, we investigate if temperature stress triggers adaptation of neuronal signaling. We used replicated laboratory evolution experiments with *Drosophila simulans* to explore the genetic and phenotypic consequences of adaptation to novel temperatures. A combination of next-generation sequencing, genetic and pharmacological manipulations enabled us to identify thermally adaptive traits that evolve at elevated temperature in controlled laboratory settings. We show an adaptive response in neuronal tissue, with lower expression of genes for dopamine synthesis, recycling and trans-synaptic signaling. This key neuronal adaptation resulted in enhanced locomotor activity in hot environments by modulating dopamine signaling with likely pleiotropic consequences. Our study provides the first direct and unambiguous evidence that environmental temperature drives adaptive changes in neuronal tissues. The discovery that dopamine function in the brain evolves in response to the environment has far-reaching impact on evolutionary biology, ecology and behavioral neuroscience.

## Main

We investigated traits that responded concordantly in ten independently evolving *Drosophila simulans* populations kept for more than 100 generations at a high-temperature selective regime (28°C/18°C, 12/12h circadian cycle). We used RNA-seq to compare gene expression of 10 hot-evolved replicates to five ancestral replicates and to five replicates evolving in a cold environment (20°C/10°C, 12/12h circadian cycle; Supplementary Figure S1).

After evolution at novel thermal regimes, the expression of 2813 genes (29% of all expressed genes, N=9283) changed significantly (FDR<0.05), showing that thermal adaptation has a strong influence on molecular traits (Supplementary Figure S2). The expression of 11% (300 genes) of the adaptive genes was significantly altered in hot-evolved replicates but unchanged in cold-evolved replicates (hot adaptive genes, Figure 1A, Supplementary Figure S2, Supplementary Table 1). Almost three quarters of the adaptive genes (74%) adapted to temperature, i.e. their expression changed in opposite directions in hot- and cold-evolved replicates, suggesting widespread thermal adaptation (temperature adaptive genes, Figure 1A, Supplementary Figure S2, Supplementary Table 1). The direction of expression changes in the remaining 15% of genes is consistent with laboratory adaptation and adaptation to cold temperature (Supplementary Figure S2, Supplementary Table 1) and these genes were not further considered.

**Figure 1.**
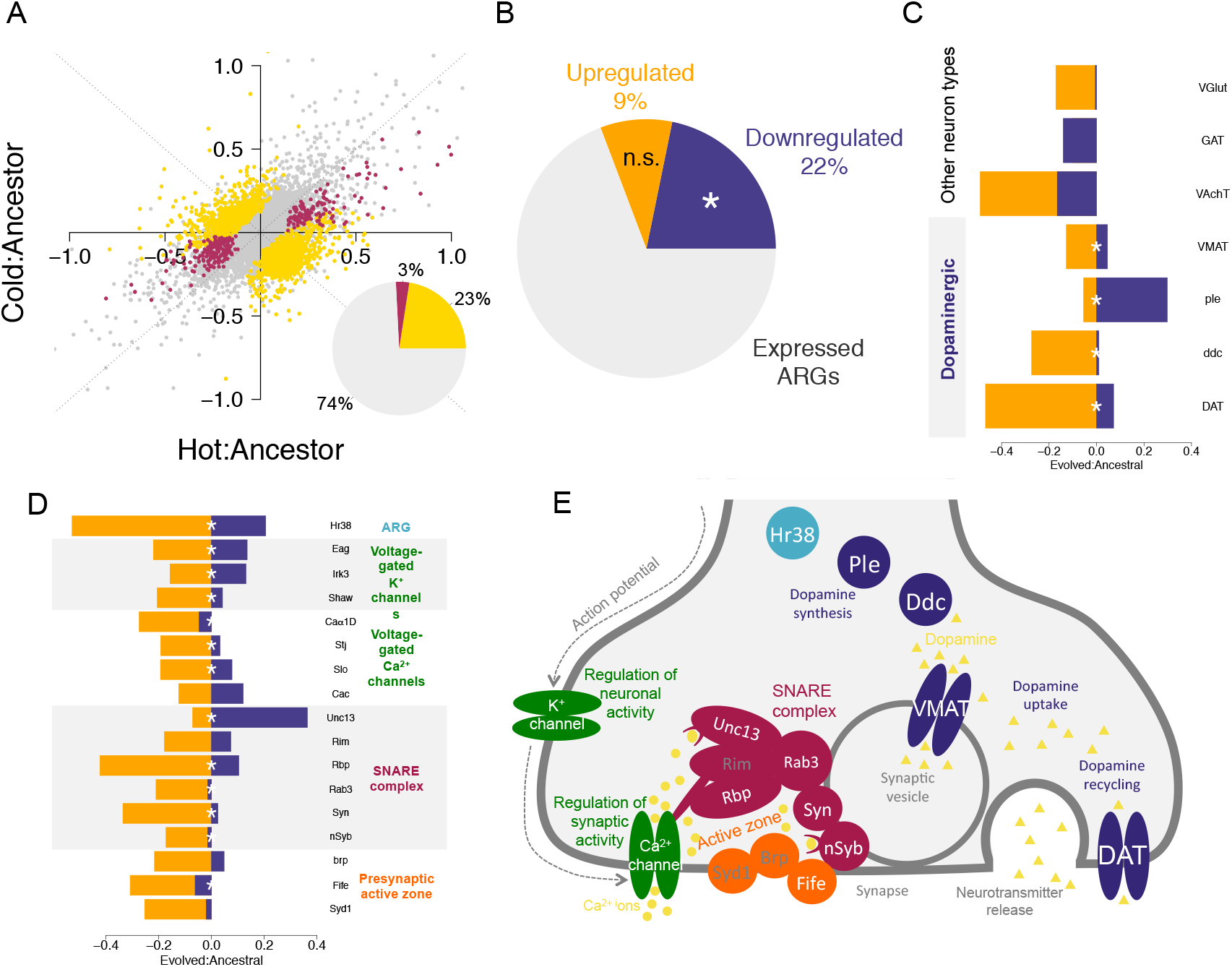
Expression of neuronal activity genes is downregulated after evolution at high temperatures. **A)** Evolution of gene expression. All expressed genes (grey) are shown. Significant changes in gene expression for hot adaptive and temperature adaptive genes are shown on log2 scale and colored red and yellow, respectively. **B)** Expressed neuronal activity-regulated genes (ARGs, N=78) are significantly enriched (**\*** p<0.05) in genes downregulated after evolution at hot temperatures. **C)** Vesicular neurotransmitter transporters, dopamine synthesis and recycling genes show significant (**\*** FDR<=0.05) temperature adaptive gene expression with hot evolved replicates (orange) decreasing, and cold evolved replicates (blue) increasing expression after evolution. **D)** and **E)** Significant temperature adaptive expression changes for essential neuronal and synaptic activity proteins. Basic neuronal function is downregulated at gene expression level in hot evolved replicates (orange) and upregulated in cold evolved replicates (blue). Significance (FDR<=0.05) is indicated with **\*** in D), and with white font in E). Functional groups of genes in D) and E) are indicated with the same color.

Genes significantly upregulated in hot-evolved replicates were enriched for several (23) sperm motility gene ontology (GO) categories, while genes downregulated in hot-evolved replicates were extensively enriched for more than 200 GO categories (Supplementary table 2). Many of these 200 categories relate to transcriptional regulation or to regulation of neuronal function and development. The enrichment for neuronal function was further supported by the significant and strong enrichment (FET odds ratio >2 and FDR<0.05) of downregulated genes expressed in head and brain tissues (Supplementary Table 3) and by a significant enrichment for neuronal anatomical categories (Supplementary Table 1). Genes encoding proteins essential for neuronal activity, presynaptic active zone and synaptic vesicle release were downregulated after adaptation to high temperature and upregulated in cold-evolved replicates (Figure 1D, Figure 1E, Supplementary Table 1). Genes downregulated after evolution at high temperature were also significantly enriched for neuronal activity-regulated genes, ARGs^9^ (p=0.005, Figure 1B, Supplementary Table 4).

Changes in the expression of vesicular neurotransmitter-specific transporters can provide insights into the type of neurons targeted by selection^10^. Remarkably, the vesicular monoamine transporter *(VMAT)* was the only transporter with a significant temperature adaptive expression change (FDR=0.05), suggesting a role of monoaminergic (dopaminergic, serotonergic, octopaminergic and tyraminergic) neurons in thermal adaptation (Figure 1C). Enrichment for anatomy ontology categories indicated signaling of dopaminergic neurons (DANs) to mushroom body neurons (MBNs) (“Calyx of adult mushroom body”, FDR=0.05; Supplementary Table 5) as a central process. Next, we specifically tested for gene expression evolution in genes dedicated to dopamine signaling. The key dopamine synthesis genes (ple and *ddc*, encoding for Tyrosine hydroxylase and Dopa decarboxylase) and the dopamine transporter gene *(DAT)* were among the temperature adaptive genes downregulated after evolution at high temperature (FDR=0.01, FDR=0.01, FDR=9.12×10^−05^; Figure 1C, Supplementary Table 1). One of the most pronounced temperature adaptive expression changes (FDR=1.07×10^−06^, Figure 1D, Supplementary Table 1) was seen for *Hr38*, which encodes a transcription factor that regulates the expression of *ple* and *ddc*^11^ and is the most strongly induced ARG in DANs and other neurons^9–10^. The dopamine receptor genes *(Dop1R1, Dop2R, Dop1R2* and *DopEcR)* showed the same temperature adaptive trend but all (except *DopEcR)* were below our threshold for detection *(DopEcR* FDR=0.26; Supplementary Figure S3). Conversely, other monoaminergic neurons did not show consistent temperature adaptive expression changes (Supplementary Figure S3). Thus, anatomy enrichment analyses in conjunction with changes of expression of neuronal marker genes implicate dopamine signaling from dopaminergic neurons (DANs) to mushroom body neurons (MBNs) as the most likely type of neurotransmission under selection.

Dopamine synthesis genes *(ple* and *ddc*) are also involved in functions unrelated to neuronal signaling, such as cuticular pigmentation^12^. There is evidence that cuticular pigmentation is selected by temperature^13^ and we found a significant, but mild decrease of whole-body melanin content in hot-evolved adults compared to the ancestral population (Figure 2A, Supplementary Table 6), with the same trend visible in flies held at 25°C (Supplementary Figure S4, Supplementary Table 6). However, we found no significant difference in thoracic cuticle pigmentation (Supplementary Figure S5, Supplementary Table 6) and the expression of genes encoding the enzymes that convert dopamine to melanin did not show temperature-or hot adaptive changes (Supplementary Figures S6 and S7, Supplementary Table 1). Because genes expressed only in neurons (*DAT, VMAT* and dopamine receptors) evolved temperature-specific changes in expression, we conclude that selection operates on a neuronal trait rather than on pigmentation.

**Figure 2.**
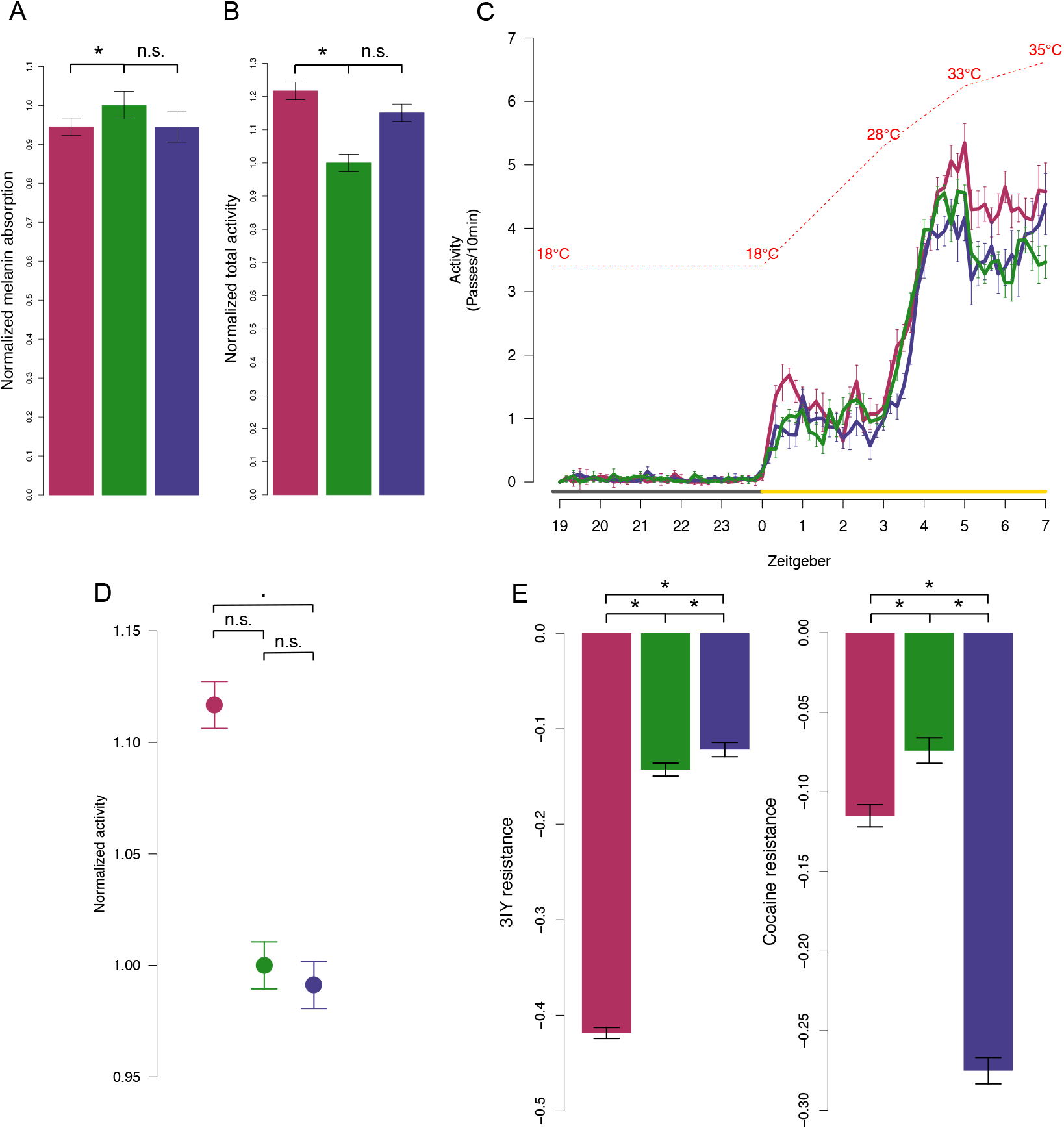
Evolved downregulation of neuronal dopamine signaling increases locomotor activity. Color code for hot-evolved (red), ancestral (green) and cold-evolved (blue) replicates and significance notation (**\*** p<0.05, ▪ p<=0.1, **n.s**. p>=0.1) is consistent in all panels. Whiskers indicate ±1SE. **A)** Melanin content. Normalized melanin-specific IR absorption is significantly reduced in hot-evolved replicates reared in the 28/18°C temperature regime. Absorption is normalized by mean estimate of ancestral replicates. **B)** Locomotor activity at 28°C. Normalized estimate of summed locomotor activity for the three evolution types. Activity is normalized by the mean estimate of ancestral replicates. **C)** Locomotor activity in a sensitized assay with increasing, high temperature. The increase in locomotor activity in hot-evolved replicates is exacerbated at very high temperatures (>33°C). In C) activity is shown as the number of passes through the light beam summed over 10 min intervals. Lines show means across replicates of each evolution type (ancestral, hot-, and cold-evolved). Temperature increase is indicated by the red interrupted line. Light condition is indicated by grey (dark) and yellow (light) lines. **D)** Total activity (normalized by ancestral mean estimate) at high temperatures (throughout the light, high-temperature period; see C)). **E)** Resistance to pharmacological perturbation of dopamine levels with 3IY and cocaine. Mean estimate of the difference in attained height between treated and control flies in a startle response assay at 28°C is shown. Significant comparisons between evolution types are indicated with asterisk.

Next, we asked whether evolution affected dopamine-regulated behaviors, such as spontaneous locomotor activity^14^. The spontaneous locomotor activity at 28°C of hot-evolved replicates was significantly higher than that of ancestral and cold-evolved flies (Figure 2B, Supplementary Table 7). The difference was exacerbated in a sensitized assay based on a temperature-ramping regime, with extreme temperatures up to 35°C (see Methods; Figure 2C and 2D, Supplementary Table 7). Because the higher activity of hot-evolved flies may be a nonspecific evolutionary response, possibly unrelated to dopamine, we pharmacologically modulated dopamine signaling in evolved and ancestral replicates. We used the startle response - a standard assay for dopamine-driven phenotypes - to test for sensitivity to cocaine (a DAT inhibitor, synaptic dopamine agonist) and 3IY (a tyrosine hydroxylase inhibitor, dopamine synthesis antagonist) at 28°C. Hot-evolved replicates were highly sensitive to 3IY, while cold-evolved replicates were highly sensitive to cocaine (Figure 2E, Supplementary Table 8). Hence, hot and cold evolved replicates exhibit low and high dopaminergic activity, respectively, validating the temperature adaptive expression of dopamine signaling genes. The evolved gene expression changes thus result in significant alterations of dopamine-mediated locomotor phenotypes in evolved *D. simulans* replicates at hot temperatures. Thus, temperature-mediated selection on dopamine genes affects neuronal behavior traits rather than pigmentation.

To examine whether the evolved gene expression changes of the hot-evolved *D. simulans* replicates are sufficient to affect dopamine-related behavior phenotypes at high temperatures, we used the GAL4-RNAi system in *Drosophila melanogaster*. We knocked down the expression of genes dedicated to dopamine synthesis, recycling and trans-synaptic signaling in all post-mitotic neurons (*elav-GAL4*), DANs (ddc-GAL4) and MBNs (247-GAL4, *mef2*) and tested the locomotor activity of the flies in the sensitized temperature ramping assay (Supplementary table 9). Knockdown of the neuronal dopamine genes significantly increased locomotor activity at high temperatures (Figures 3A, 3B; Supplementary Figures S8, S9, S10, Supplementary Table 10). Typically, only weaker, long hairpin knockdowns^15^ resulted in highly active flies; strong knockdowns using short hairpin RNAi for *ple* or neuronal dopamine-deficient *ple* mutant lines^16^ strongly impaired locomotor activity independent of temperature (Supplementary Figure S11), indicating a fine-tuning of the adaptive phenotype. Targeting RNAi in DANs and MBNs separately revealed the dopamine-related function of *ple, ddc, VMAT, DAT* in DANs and *Dop1R1* in MBNs (Figure 3B). This uncovered the potential autoregulatory roles for *Dop1R1, Dop2R* and *DopEcR* in DANs. This suggests that dopamine-dependent locomotor activity is regulated by a positive, temperature-dependent feedback loop via DAN-expressed dopamine autoreceptors, as suggested for learning and memory^10,17^. Conversely, pannier-GAL4-driven RNAi of *ple, ddc, VMAT* and *DAT* expression during thoracic cuticle formation showed that dopamine synthesis is required for thoracic cuticular pigmentation, while synaptic dopamine recycling is not (Figure 3C). Together, this provides direct functional evidence of the genes and the neuronal cell types mediating the evolved temperature response.

**Figure 3.**
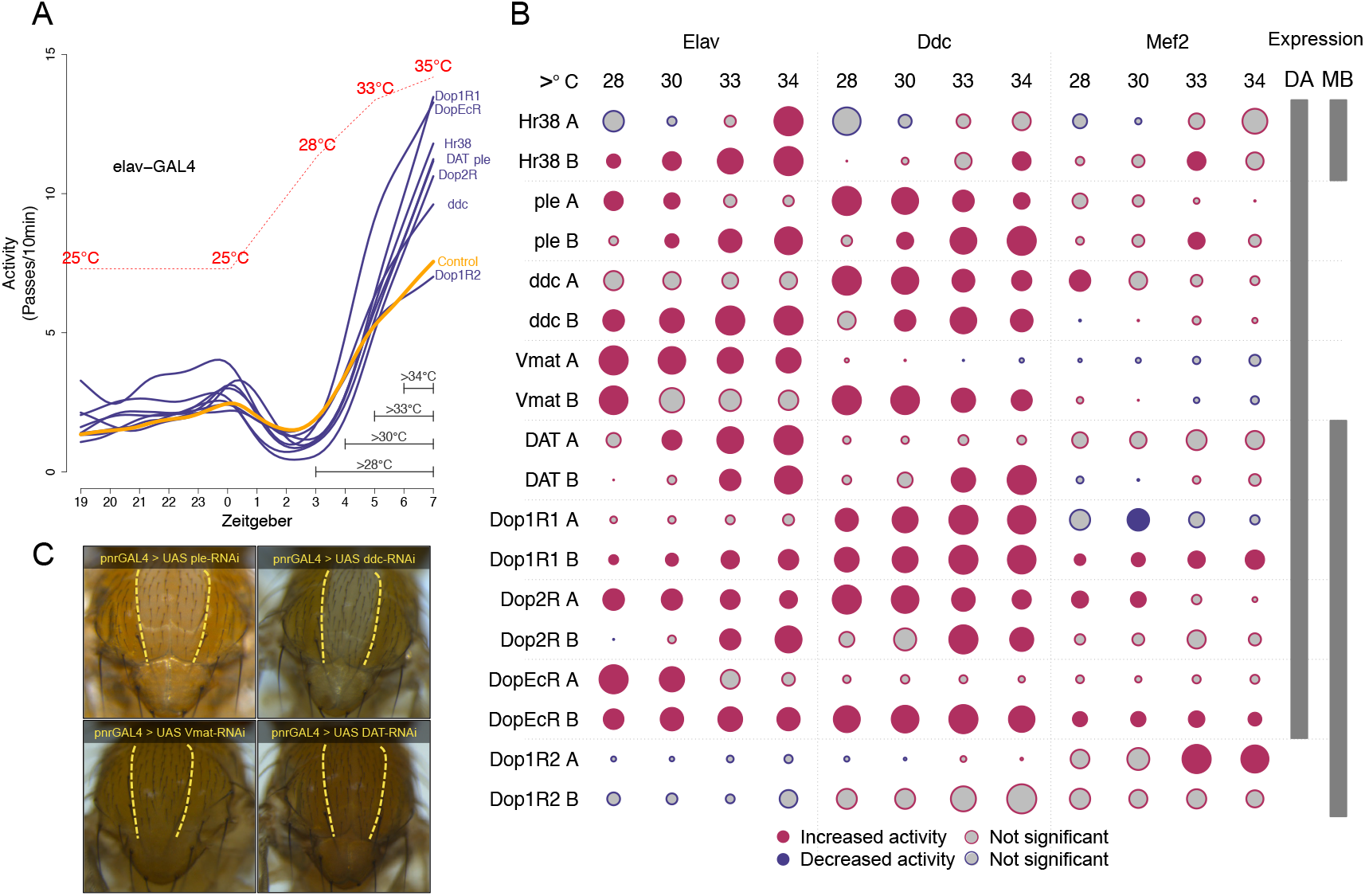
Downregulation of neuronal dopamine pathway is sufficient for the increase in locomotor activity at high temperatures. **A)** Locomotor activity after dopamine gene knockdown in post-mitotic neurons. Figure shows spline fit through mean activity across individuals. The strong increase in activity at 33°C matches the pattern of hot-evolved replicates in Figure 2C). **B)** Locomotor activity at different high-temperature ranges after dopamine gene knockdown. Activity at temperatures >28°C, >30°C, >33°C and >34°C are shown for different GAL4 (columns) and UAS-RNAi (row) crosses. Grey horizontal whiskers in panel A) indicate the tested temperature ranges. Each gene knockdown was assessed in two different UAS-RNAi; lines (A and B, see Supplementary Table 9). The diameter of the circle indicates the normalized effect size, while the color shows the direction (higher activity=red lower activity=blue) and the significance of change (not significant=grey, p<0.05=colored) in activity compared to the control line. Grey bars on the right side indicate the expected localization of expression for a given gene. **C)** pnr-GAL4 driven RNAi of dopamine synthesis and recycling genes, *ple* and *ddc* RNAi in cuticle affects pigmentation while DAT and VMAT RNAi do not. Yellow interrupted lines indicate localization (borders) of pnr-GAL4 expression.

In summary, we present multiple, independent lines of evidence that synaptic transmission and neuronal dopamine signaling evolves during thermal adaptation of *Drosophila simulans*. The lower expression of dopamine-related genes in the brains of evolved flies allows higher locomotor activity at increased temperatures. We propose that the high temperature stimulates DAN hyperactivity in ancestral flies, decreasing their locomotor ability and consequently their fitness. The evolution of lower dopamine signaling alleviates the fitness cost of the high temperature. This notion is consistent with dopamine-related thermal sensitivity, where low-dopamine flies prefer higher temperatures and vice versa^18^. Different individual DAN circuits form the basis of neuronal dopamine pleiotropy, regulating sleep, arousal, aggression, courtship and valence to sensory perceptions in learning and memory^19^. We therefore assume that spontaneous locomotor activity is only one phenotypic consequence of the evolved dopamine signaling and other dopamine-related behavioral traits remain to be identified.

How do these results from laboratory evolution relate to natural populations? There are latitudinal allele frequency clines of genes encoding neuronally expressed dopamine receptors *(Dop1R1, Dop2R, DopEcR)* and transporters (DAT and VMAT) in natural populations of *Drosophila* and *Anopheles*^20–24^ (Supplementary Table 11). This strongly argues that the evolution of temperature-dependent neuronal dopamine signaling is not a laboratory artefact but a common response of insects to selection at high temperatures in nature. We propose that the evolutionary response of dopamine signaling to elevated temperatures is an important step to mitigate the deleterious impact of temperature change. Assuming that the pleiotropic architecture differs between species, we anticipate an increase in phenotypic divergence across species for different traits regulated by dopamine, such as pigmentation, locomotion, courtship and other behaviors^25–27^.

## Methods

### Evolution experiment

202 isofemale lines were combined 15 times to seed each of the 15 replicated populations. Of those, 10 replicates were maintained at the high temperature regime 12h/12h 28°C/18°C and 12h/12h light/dark circadian cycle for 100 generations (hot evolved replicates; as described in ^28^) while 5 replicates (cold-evolved replicates) were maintained at the low temperature regime 12h/12h 20°C/10°C and 12h/12h light/dark for over 50 generations (Supplementary Figure S1). Population size was maintained each generation between 1000 and 1250 adult individuals per replicate.

### RNA-seq common garden experiment

RNA-seq common garden was performed after 103 generations at high temperature regime (or 52 generations in low temperature regime, Supplementary Figure S1). We pooled around 180 founder isofemale lines (some were lost over the years) 5 times to seed 5 replicates of the reconstituted ancestral population. Both evolved and ancestral replicates (10 hot evolved, 5 cold evolved and 5 ancestral replicates) were reared for at least 2 generations in the same conditions. Whole 50 5-day-old males were flash-frozen in liquid nitrogen and stored at −80°C until RNA extraction.

### RNA-extraction and library preparation

Flies were homogenized in Qiazol (Qiagen, Hilden, Germany) and total RNA was extracted from the homogenate using the Qiagen RNeasy Universal Plus Mini kit with DNase I treatment. Barcoded and stranded mRNA libraries were prepared using the TruSeq stranded mRNA Library Prep Kit for Neoprep (Illumina, San Diego, USA). Neoprep runs were performed using software version 1.1.0.8 and protocol version 1.1.7.6. starting with 100ng of total RNA. All samples were randomized across library cards, and library cards with the same lot number were used for all Neoprep runs to avoid batch effects. Barcoded libraries were quantified using the Qubit DNA HS assay kit and pooled in equimolar amounts. Reads with a length of 50bp were sequenced on an Illumina HiSeq 2500 platform.

### Differential expression analysis

All RNA-seq reads were trimmed based on quality (−min_quality 20) with ReadTools^29^ and mapped to the *D. simulans* genome^30^ using GSNAP^31^ with enabled split read mapping (-N 1) and relaxed mismatch parameter (-m 0.08) to account for polymorphisms. Reads were counted based on the *D. simulans* annotation^30^ using RSubread^32^. Differential expression analysis between the evolved and ancestral replicates was performed in the EdgeR framework^33^ after filtering 30% of genes with the lowest expression. To test for differential gene expression, we modeled evolution type (“ancestral”, “hot-evolved”, “cold-evolved”) and anchored ancestral expression as the intercept (counts ~ 0 + evolution). Different evolution types were assigned based on criteria in Supplementary Table 12. All p-values were FDR corrected for multiple testing with the significance cutoff FDR < 0.05.

### Enrichment tests

Gene ontology enrichment was performed using GOrilla^34^ with all expressed genes (N=9283, Supplementary Table 1) as the background set. Tissue and anatomy ontology enrichment were calculated using Fisher’s exact test and were based on FlyAtlas2 expression data^35^ and the Flybase anatomy ontology database from FlyBase Consortium^36^ (obtained from Victor Strelets, Supplementary File 1). Where required, significance values were adjusted for multiple testing.

### Locomotor activity assays

Spontaneous locomotor activity was measured for ancestral and evolved replicates (hot-evolved generation F140 and cold-evolved generation F70; Supplementary Figure S1) using Drosophila Activity Monitor System (DAMS, TriKinetics, USA). After two generations at common garden conditions in the high-temperature regime activity was recorded either under the native high-temperature regime assay (28/18°C) over 6 days or at a sensitized high-temperature ramping assay with gradual temperature increase from 18°C to 35°C over 7 hours (as indicated in Figure 2C). To test for consistency across independently evolving replicates we compared MODEL1 y_*i*_, = X_*E*_β + Zμ_*r,m*_ + *ε_i_* and MODEL2 y_*i*_, = Zμ_*r,m*_ + ε_*i*_ fits within an ANOVA framework, where y is summed activity for each individual (*i*) in each replicate (*r*) of a given evolution type (*E*) in a particular measurement day (*m*) using lmer function in lme4 R package (fixed and random effects are indicated in upper and lower case, also in the following models). Pairwise differences were tested using Tukey’s post-hoc test with the TukeyHSD function in lsmeans R package (Supplementary Table 7).

### Drug sensitivity

To measure sensitivity to perturbation of dopamine signaling we employed the startle response assay after pharmacological treatment with cocaine (Cocaine, COC-156-FB, Lipomed AG, Austria) or 3IY (3-Iodo-L-tyrosine, I8250, Sigma Aldrich, USA). Flies were reared for two generations in the high-temperature regime common garden experiment (Supplementary Figure S1). We treated 50 flies with 2μg of cocaine dissolved in ethanol and vaporized using a “fly crack pipe” (1 minute exposure) constructed as described in^37^. For 3IY treatment 50 flies were treated with 2 mg 3IY dissolved in 2% yeast and 5% sucrose solution (soaked filter paper) for 3 days. Flies were transferred to a 100ml glass graduated cylinder and after a short (~30 s) acclimatization period forced to the bottom of the cylinder by firmly tapping it on the bench. Control and treated flies in each replicate were assayed at the same time. The startle response was recorded with a Samsung Galaxy S6 camera (high definition resolution, 60 fps capture, white balance 38000 and auto-focus) and measured as the average attained height of the 50 flies 30 seconds after being startled. Height was assessed using the volume marks on the cylinder. The sensitivity to chemicals was measured as the difference between startle response of the treated flies and the untreated flies reared under the same conditions. The interaction between evolution type and drug effect was assessed with the AIC goodness of fit of MODEL1 y_*i*_ = X_*DxE*_β + Zμ_*r,m*_ + ε_*i*_ compared to reduced MODEL2 y_*i*_ = Zμ_*r,m*_ + ε_*i*_ where y is the attained height 30 sec after startle response for an individual fly *i* of a given replicate (*r*) and measurement (*m*) of evolution type (*E*) in a control or drug treatment (*D*). The differences in sensitivity to the drug between evolution types were estimated as an interaction between *E* and *D* (Supplementary Table 8).

### Pigmentation and melanin content

To assay pigmentation and melanin changes, flies were reared at constant 25 °C common garden and very low egg density (150 eggs/bottle) or 28/18 °C and low egg density (400 eggs/bottle; Supplementary Figure S1). Flies from 25 °C common garden were mounted on a cover slide submerged in 70% ethanol and thorax images were taken the same day using the same imaging parameters and illumination. Pixel color was quantified for images of ancestral and evolved replicates, as well as *yellow* (melanin-deficient) and *ebony* (highly pigmented) mutant *D. simulans* strains as extreme pigmentation reference points (Supplementary Table 9). Between-group analysis with subsequent leave-one-out predictive analysis and data projection^38^ was used to quantify pigmentation change in evolved replicates. To determine relative melanin content, 200 males from 28/18°C and 25°C common garden (ancestral, evolved and yellow mutants) were vacuum-dried at 80°C for 72 h and pulverized with a porcelain mortar into fine powder (~5-7 mg of powder per sample was obtained) and homogenates were analyzed by infrared (IR) spectroscopy. To obtain IR spectra of a thin uniform layer, 3-5 mg of homogenized powder was pressured on a diamond ATR plate using self-levelling sapphire anvil of a Cary 660 Fourier transform infrared (FTIR) spectrometer (Agilent Technologies) coupled with Golden Gate single-reflection diamond Attenuated Total Reflectance (ATR) accessory (Specac). Absorption spectra were recorded in a mid-infrared region (4000-400 cm^−1^) at a nominal resolution of 4 cm^−1^ (at RT; 23 ± 1 °C). Two replicate spectra (50 scans/spectrum) of each sample were recorded using different aliquots. Molecular vibrations observed in IR spectra were assigned using electronic spectral libraries (NIST, Agilent Technologies, IMB Jena Image Library of Biological Macromolecules) and published spectral data (see Supplementary Methods 1). Relative melanin content was determined from the IR absorbance for determined melanin-specific wavenumbers with melanin absorption index (MAI) >1 (MAI maxima at 2925, 2853, 1743, 1153, 1077 and 1023 cm^−1^; see Supplementary Methods). Full and reduced general mixed models were fitted to the melanin absorbance data (lme4 R package; MODEL1 y_*i*_ = X_E_β + Zμ_r,m_ + ε_i_ and MODEL2 y= Zμ_r,m_ + ε_i_, (where y is absorbance value for each individual *i*, in each replicate *r*, measurement *m*, of evolution type *E*) to test for replicate consistency (Supplementary Table 6). The best model fit was chosen based on AIC score within an ANOVA framework. Differences between evolution types were tested using Tukey’s post-hoc test (lsmeans R package).

### GAL4-UAS knockdown assays

The *D. melanogaster* strains used for genetic knockdown are given in Supplementary Table 9. Mini-*white*^+^ GAL4 transgenes and *white* genetic backgrounds have been suggested to alter several behavioral traits in males and to affect dopamine levels^39–41^. Therefore, we crossed RNAi line virgins to GAL4 line males, making the assayed F1 males exhibit wild-type eye colors for Bloomington and VDRC/KK RNAi lines, and very dark eye color for VDRC/GD lines. Flies were raised at 25°C in three replicates for the activity assay after neuronal knockdown. Locomotion was assayed for 20-40 males per line. Flies were aged 3-6 days and assayed in a sensitized high-temperature assay (see “Locomotor activity assays”) at 25°C during the dark part of the cycle. All crosses were assayed in parallel, including controls, in multiple measurements. We used Dunnet adjusted pairwise comparison between KD crosses and appropriate controls to account for comparisons with the same control. Comparisons were performed using mean activity estimated after fitting a model y_*i*_, = X_*L*_β + Zμ_*r,m*_ + ε_*i*_ where y is summed activity measured at a temperature above a threshold (see Figure 3A) for individual fly *i* from GAL4xUAS-RNAi line cross L, replicate *r* and measurement *m* (Supplementary Table 2).

A single cross was produced for the cuticular knockdown with *pnr*-GAL4. Flies were mounted and imaged following the same protocol as for the pigmentation assay. Due to the severity of the effects on pigmentation, it was possible to assess cuticle discoloration visually by assigning a binary value (“changed” or “unchanged”).

### Data availability

All raw RNA-seq data will be uploaded to NCBI project XXX and available upon publication. All raw data and full code used to produce results (raw RNA-seq count tables, raw locomotor activity data, startle response data, pigmentation images, raw IR spectra) will be uploaded and available for download on DataDryad upon publication.

## Acknowledgments

We thank the Birman, Knoblich, and Keleman labs, and VDRC and BDSC for *D. melanogaster* lines, the FlyBase Consortium for the anatomy ontology file, Miljenko Jakšić, Ivan Tomić, Marko Jakšić and Jay Hirsh for the advice and equipment needed to construct the “fly crack pipe” and R. Mazzucco and M. Dolezal for advice on the statistical analysis, Graham Tebb, Andy Clark and Yasir Ahmed-Braimah for comments on the manuscript. This study was funded by ERC, FWF and Marie Curie Actions.

## Author contributions

A.M.J., K.A.S. and C.S. conceived the study and interpreted the data.

C.S. designed, initiated and maintained the evolution experiment.

A.M.J., V.N., N.B., F.M., K.A.O. carried out the RNA-seq common garden experiment

L.S. and A.M.J. conceived the spectroscopic melanin content measurement method, and analyzed and interpreted the data.

S.K.H. contributed to RNA-seq data analysis and set up common garden experiments.

A.M.J. and K.A.S. designed RNAi knockdown and mutant knockout activity experiments.

A.M.J., J.K. and K.A.S. performed RNAi knockdown and mutant knockout activity experiments.

A.M.J. designed and carried out the rest of the experiments and analyzed the data. V.N. prepared all RNA-seq libraries and supervised the maintenance of the evolution experiment.

K.A.S. and C.S. co-supervised AMJ.

A.M.J., K.A.S. and C.S. wrote the manuscript.

## Competing interest

Authors declare no competing interest.

## Materials and Correspondence

Correspondence and material requests should be addressed to Christian Schlötterrer.

