## Supplementary Figures for "Neuronal function and dopamine signaling evolve at high temperature in *Drosophila*"

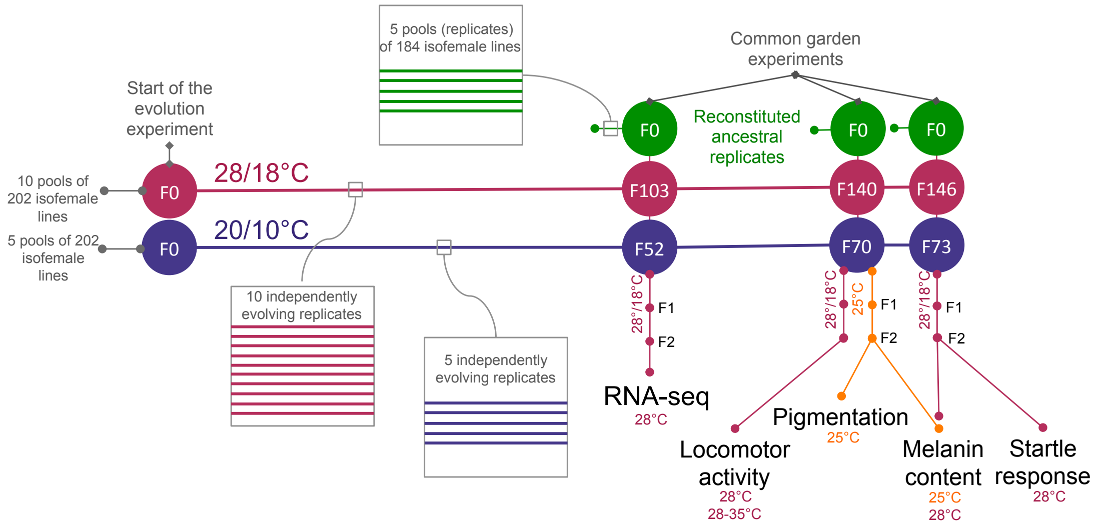


**Figure S1 Experimental timeline.** Fifteen independently evolving replicate populations (5 adapting to cold (blue) and 10 to hot (red) temperature regime) were set up by pooling 202 isofemale lines. Common garden experiments were set up at three time points. For each common garden experiment, ancestral replicate populations were reconstituted (green) from the same isofemale lines used at the beginning of the evolution experiment. Rearing temperature of a common garden is indicated vertically, whereas temperature at the time-point of sampling/assay is indicated below the assay label. All flies in the common garden experiments were reared at high temperature regime (28/18°C), except flies used for pigmentation assay and additional melanin content assay, which were reared at constant 25°C to control for potential phenotypic variance caused by thermal developmental plasticity of melanin synthesis.

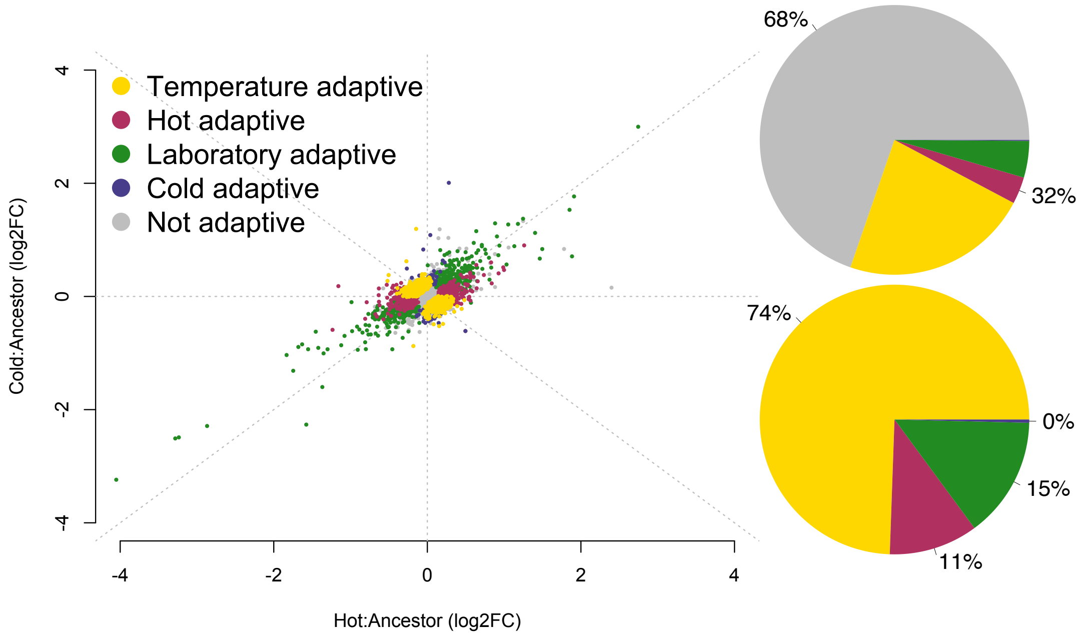


**Figure S2 Gene expression change for all expressed genes.** Scatterplot and pie charts share the color code. Upper pie chart shows proportion of all expressed genes, whereas lower pie chart highlights the proportions of the evolved, adaptive genes.

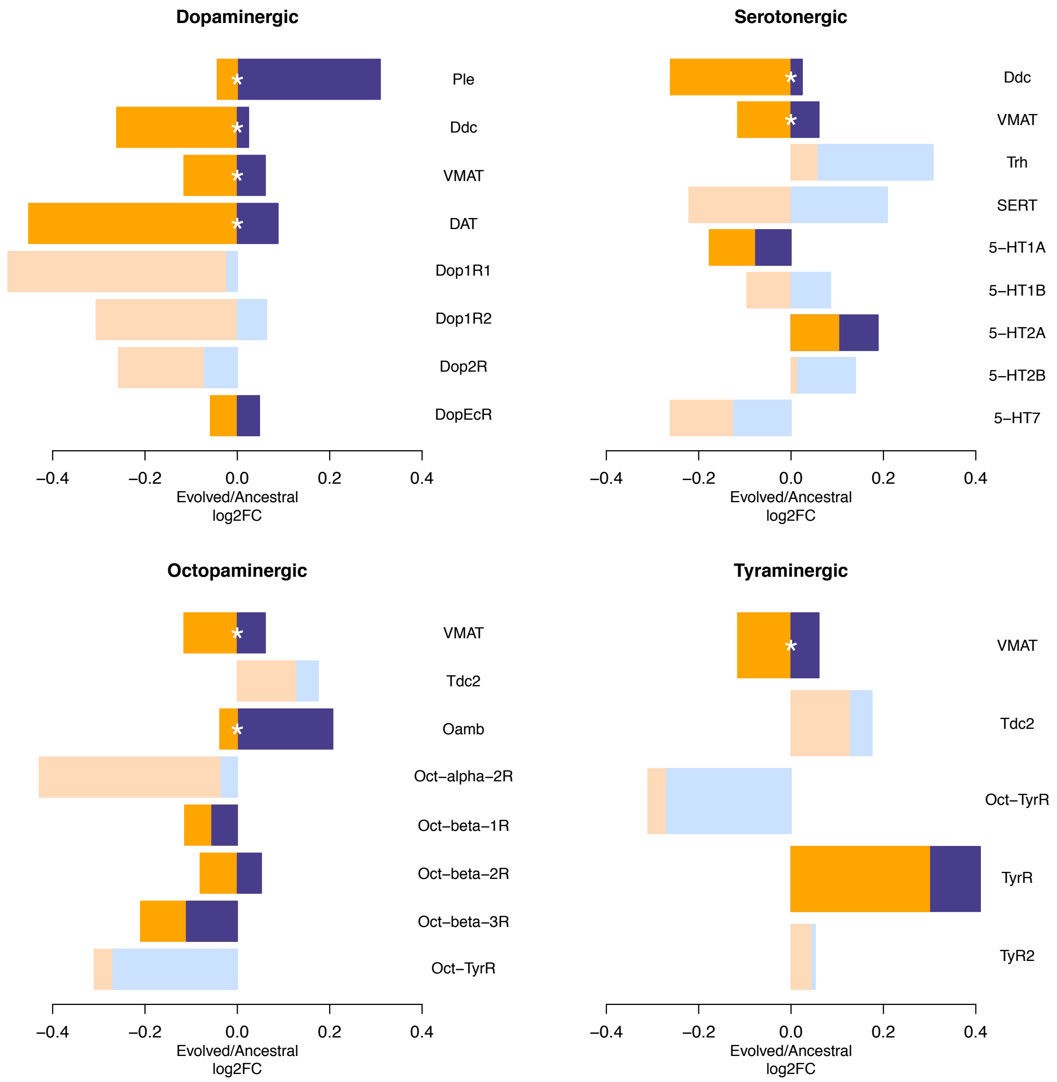


**Figure S3 Gene expression of monoaminergic neuronal marker genes.** Four panels show log2 fold change in gene expression between ancestral and evolved replicates (orange=hot evolved, blue=cold evolved) for marker genes expressed in four types of monoaminergic (VMAT expressing) neurons. Genes that did not reach our lower expression threshold are indicated in lighter colors. Significant temperature adaptive gene expression change is indicated by white asterisk.

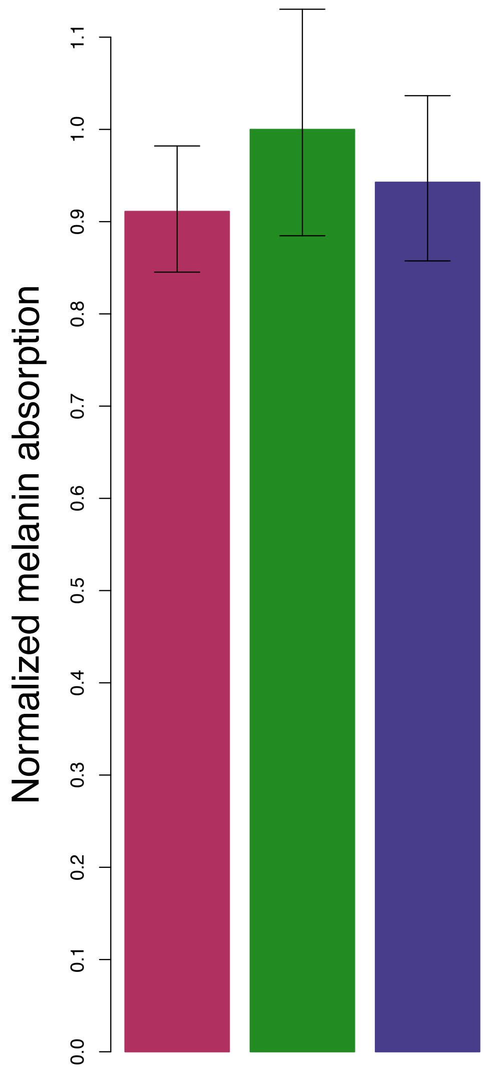


**Figure S4 Melanin content is not driven by thermal plasticity.** Melanin synthesis in *Drosophila* is thermally plastic and occurs during the last days of pupal development and first 24 h post-eclosion. To control for the influence of daily temperature fluctuations that may occur at different melanin synthesis time-points across populations (for example if pupation times differ across populations) we reared flies at constant 25°C. This ensured that every population experienced the same developmental temperature over the full melanin synthesis developmental period. Measured melanin content at 25°C was consistent with measurements at 28/18 °C (see Figure 2C in main text), although the same statistical significance was not attained (see Supplementary Table 6).

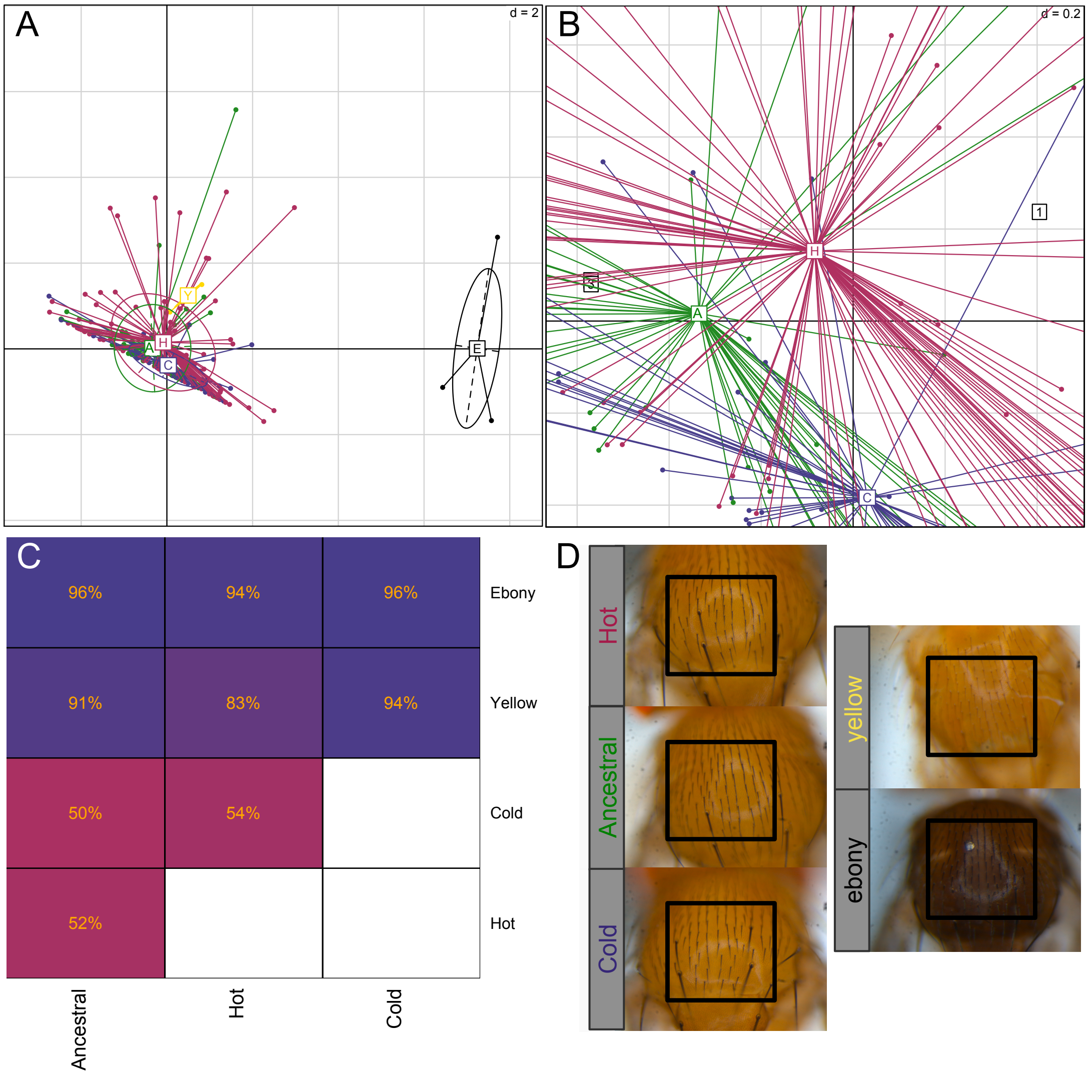


**Figure S5 Thorax pigmentation**. A) Principal component analysis for thorax pigmentation for ancestral (green, A), hot evolved (red, H), cold evolved (blue, C) populations and *yellow* (yellow, Y) and *ebony* (black, E) mutants indicates that hot evolved populations are lighter than the ancestral and cold evolved flies. **B)** Zoom-in to centroids of panel A). **C)** Percentage of correct sorting of permuted data into population groups in pairwise comparisons. Lower percentage (and lighter, pink color) indicates the two populations are more difficult to discern apart whereas higher percentage (more blue) indicates higher (clearer) divergence between populations. Only ebony mutants are significantly different than the other populations. Hot evolved populations are however more similar to yellow mutants than the other two populations.  **D)** Example of the photos and thorax area (black squares) for which color was quantified.

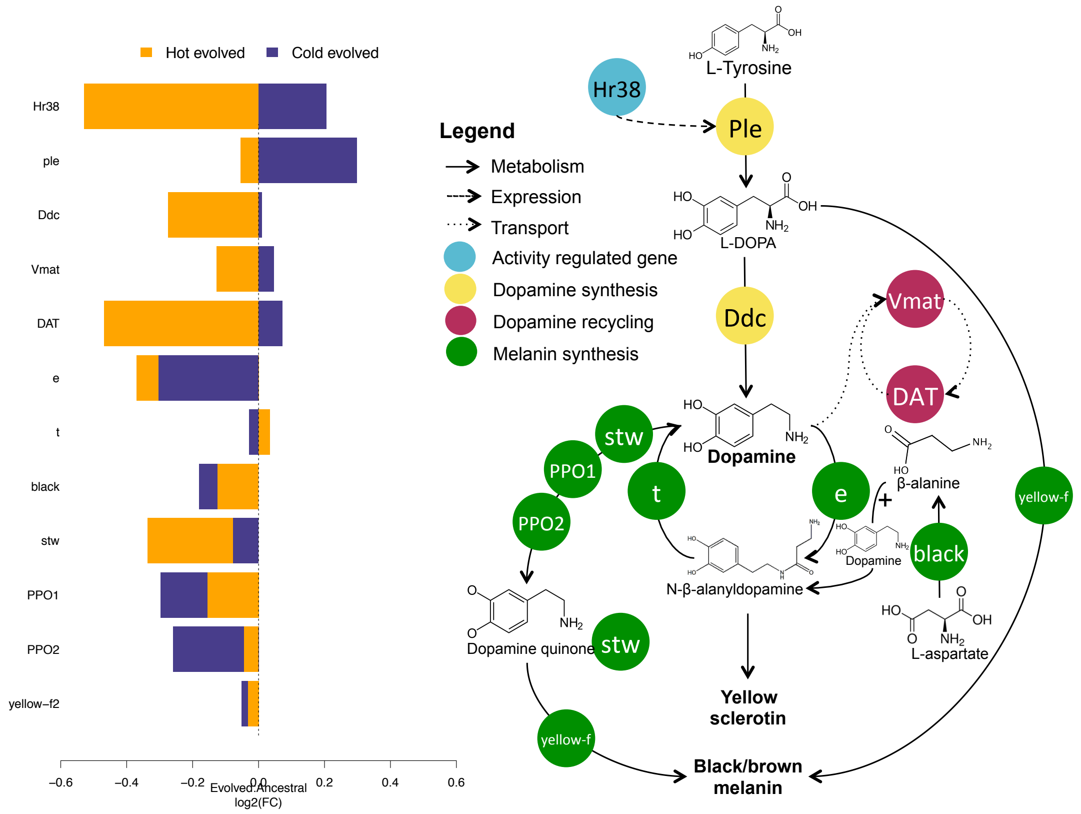


**Figure S6 Dopamine- and melanin pathway genes.** Expression changes after evolution in hot and cold temperatures are shown in the left panel, while a schematic of the pathway is shown on the right. Melanin synthesis genes downstream of dopamine do not show temperature or hot adaptive change in gene expression.

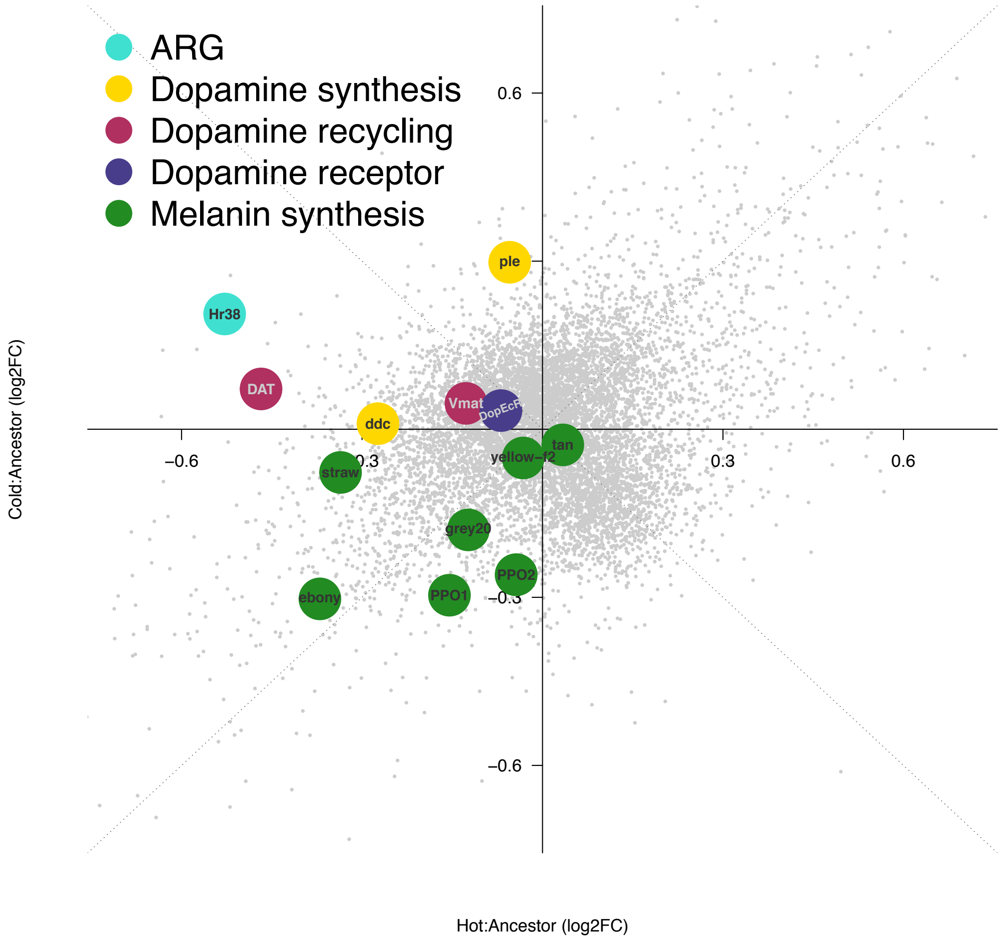


**Figure S7 Evolution of dopamine genes.** Gene expression changes for expressed dopamine- and melanin pathway genes after evolution at high and low temperatures in relation to gene expression changes of other expressed genes (grey).

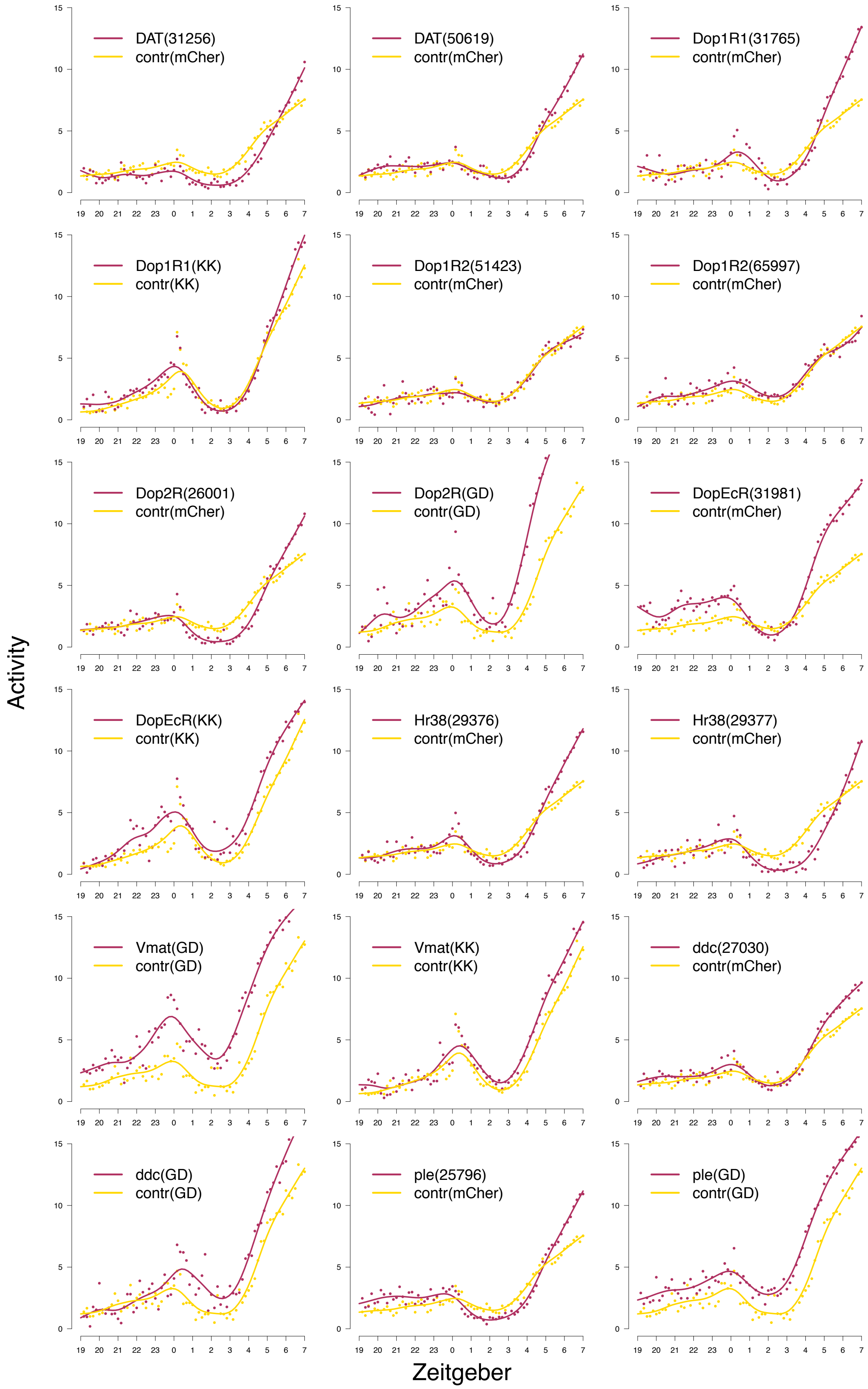


**Figure S8 Elav-GAL4>RNAi activity.** Locomotor activity at sensitized high-temperature regime for post-mitotic neuron knockdown of dopamine genes.

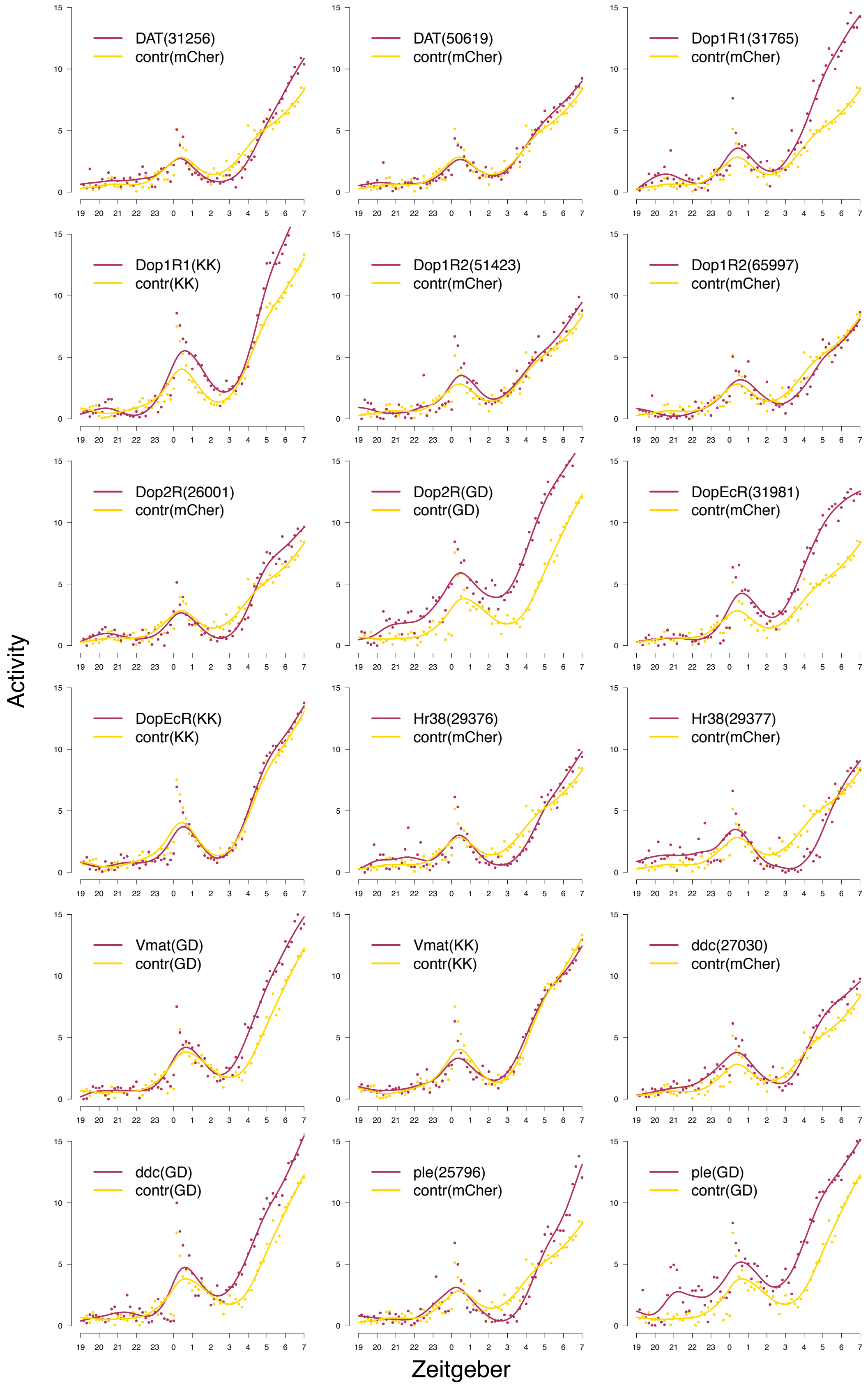


**Figure S9 Ddc-GAL4>RNAi activity.** Locomotor activity at sensitized high-temperature regime for dopaminergic neuron knockdown of dopamine genes.

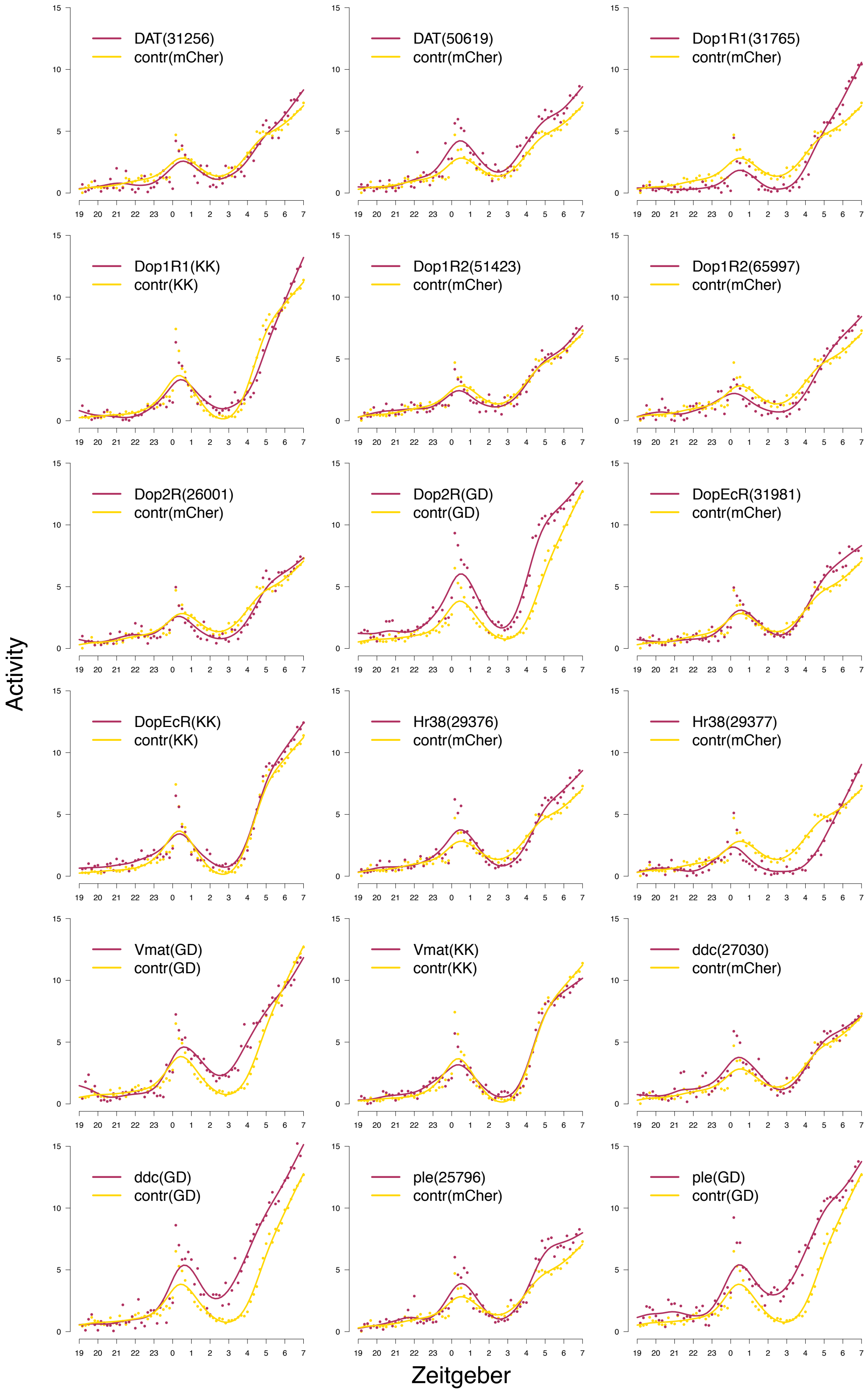


**Figure S10 Mef2-GAL4>RNAi activity.** Locomotor activity at sensitized high-temperature regime for mushroom body neuron knockdown of dopamine genes.

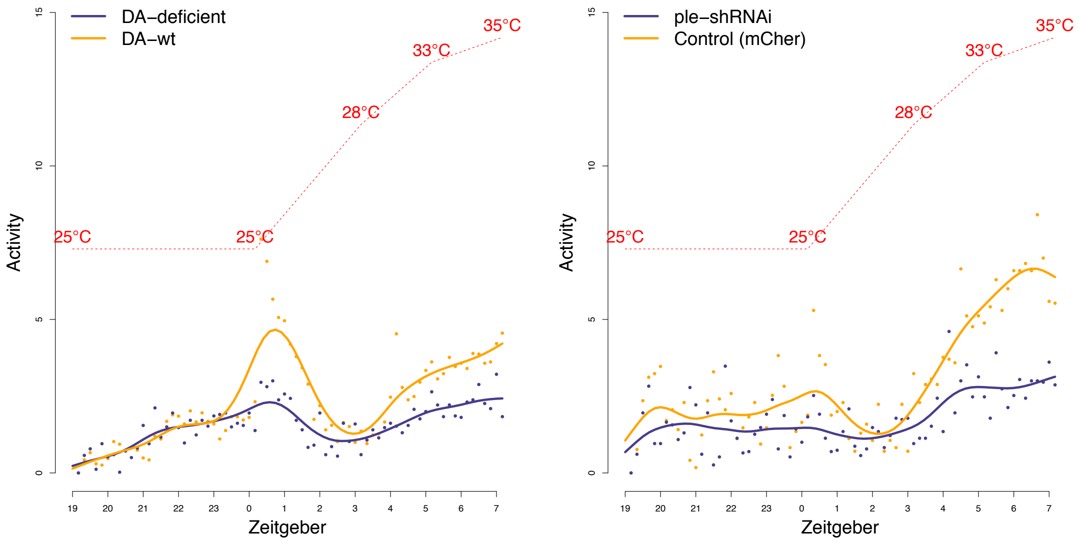


**Figure S11 Activity in strongly neuronally DA-deficient flies.** Knockout or strong knockdown of neuronal dopamine synthesis strongly reduces locomotor activity. Left panel shows locomotor activity for $DTH^{FS+/-}$ neuronal-DA-deficient *ple* rescue mutant (and wild type rescue control) while left panel shows activity for stronger short-hairpin RNAi knock-down of *ple* (and control). Line shows the spline fit for summed passes through the IR beam in 10 min intervals (points).
